# Force-biased nuclear import sets nuclear-cytoplasmic volumetric coupling by osmosis

**DOI:** 10.1101/2022.06.07.494975

**Authors:** Fabrizio A. Pennacchio, Alessandro Poli, Francesca Michela Pramotton, Stefania Lavore, Ilaria Rancati, Mario Cinquanta, Daan Vorselen, Elisabetta Prina, Orso Maria Romano, Aldo Ferrari, Matthieu Piel, Marco Cosentino Lagomarsino, Paolo Maiuri

## Abstract

In eukaryotes, cytoplasmic and nuclear volumes are tightly regulated to ensure proper cell homeostasis. However, the detailed mechanisms underlying nucleus-cytoplasm volumetric coupling remain unknown. Recent evidence supports a primary role of osmotic mechanisms in determining a tight link between nuclear and cytoplasmic volume, but this hypothesis remains largely untested in mammalian cells. We approach the question in single cultured adhering human cells, by jointly measuring cytoplasmic and nuclear volumes, in real time and across cell cycles. Surprisingly, we find that cytoplasmic and nuclear volumes follow different average growth laws: while the cytoplasm grows exponentially, the nucleus grows linearly. Moreover, by combining several experimental perturbations and analyzing a mathematical model including osmotic effects and tension, we conclude that the mechanical forces exerted by the cytoskeleton on the nuclear envelope can strongly affect nucleus-cytoplasm volumetric coupling by biasing nuclear import. Our results unveil how osmo-mechanical equilibrium regulates nuclear size in mammalian cells.

**One-Sentence Summary:** Cytoskeletal forces exerted on the nuclear envelope impact on nuclear volume through modulation of force-coupled nucleo-cytoplasmic transport, affecting osmosis.

## Main Text

In all living systems, size control of cells and intracellular organelles is essential for the optimal regulation of several biological functions (*1–3*). In multicellular organisms, cell and organelle size is a characteristic feature of a given cell type, and both cell-to-cell and mean variations are often associated to pathological conditions such as cancer or aging (*4, 5*). The size of the nucleus, the largest cellular organelle, generally scales linearly with cell size. In yeast, nuclear to cell volumetric ratio (karyoplasmic ratio or NC ratio) is roughly constant along the entire cell-cycle (*6–8*). However, for mammalian cells, the detailed mechanisms underlying nucleus-cytoplasm volume coupling are still mostly unclear (*9*).

To address this problem, we developed a technique, which we named nuclear fluorescence exclusion microscopy (N2FXm), for measuring jointly, and in real time, both nuclear and cytoplasmic volumes of living cells. This method was conceived as an evolution of fluorescence exclusion microscopy (FXm), an imaging-based technique that allows measuring cellular volumes using low-magnification objectives (*10–12*). In standard FXm, cells are injected in a microfluidic chamber of known height filled with a fluorescent dye, in our case high molecular weight RED-dextran, which is not internalized by cells. For any object in the chamber, the corresponding drop in dextran fluorescence is linearly proportional to its volume. Consequently, dextran fluorescence can be scaled linearly between two points of known height: zero, where the chamber is empty, and the maximum, the known chamber height, in correspondence of the pillars sustaining the chamber roof. This calibration allows us to define the optical thickness of cells of arbitrary shape in the chamber and, accordingly, cell volume can be precisely computed by integrating it over the segmented cell area (see materials and methods, Fig. S1A-B). N2FXm extends the FXm technique to nuclear volume measurements. Cell nuclei were negatively stained with the ectopic expression of a green fluorescent protein coupled with a nuclear export signal (GFP-NES), which localizes in the entire cytoplasm except the nucleus, which is also marked with H2B-BFP (see materials and methods, Fig. 1A-B and S1C-E). In this doubly-labeled system, and in presence of the extracellular dye, a second calibration is introduced and the cytoplasmic GFP fluorescent signal is scaled accordingly to cell optical thickness. The two calibration steps generate calibrated images for both cell and cytoplasm, obtained from dextran and GFP fluorescent signals, respectively. Since the latter is scaled according to the former, intensity profiles from the two images differ only in correspondence of the nucleus, where the difference of the two is proportional to the nuclear height (Fig. S1C). Finally, to measure nuclear volume, the calibrated GFP signal is integrated within the segmented nuclear region, subtracting the resulting value from the cell volume evaluated over the same domain (see materials and methods, Fig. S1-E and Supporting video S1). Cytoplasmic volume is obtained by subtracting the nuclear volume from the entire cell volume and the NC ratio as the ratio between nuclear and cytoplasmic volumes. To test our method, we used N2FXm to measure the volumes of spherical polymeric μ-particles (DAAM particles (*13*), see material and methods) internalized by RPE1 cells. N2FXm measurements were in good agreement with the volumes estimated with a geometrical measurement (see materials and methods and Fig. SG-I, linear fit coefficients of SI: slope=0.94, Pearson correlation coefficient=0.89, R^2^=0.78). Moreover, the distribution of the volume ratio estimated by N2FXm and geometrical measurements was compatible with a normal distribution centered on 1, excluding possible systematic errors in our procedure (Fig S1H). We also compared nuclear volume distribution of RPE1 cells measured with both N2FXm and 3D confocal reconstruction (see materials and methods) obtaining similar results (Fig. S1F). Finally, we assessed if the ectopic expression of GFP-NES or H2B-BFP perturbed cell or nuclear dimensions and we did not find any meaningful differences (Figure S1J-K). To summarize, the main advantages of N2FXm are: 1) both cell and nuclear volumes are obtained from a single plane illumination, avoiding 3D reconstruction; 2) the technique works with small magnification and low numerical aperture objectives, enabling the simultaneous record of many cells in the same field of view; 3) the imaging needs low illumination, reducing phototoxicity and making the technique suitable for long-term experiments (see supporting video S1).

**Fig. 1.**
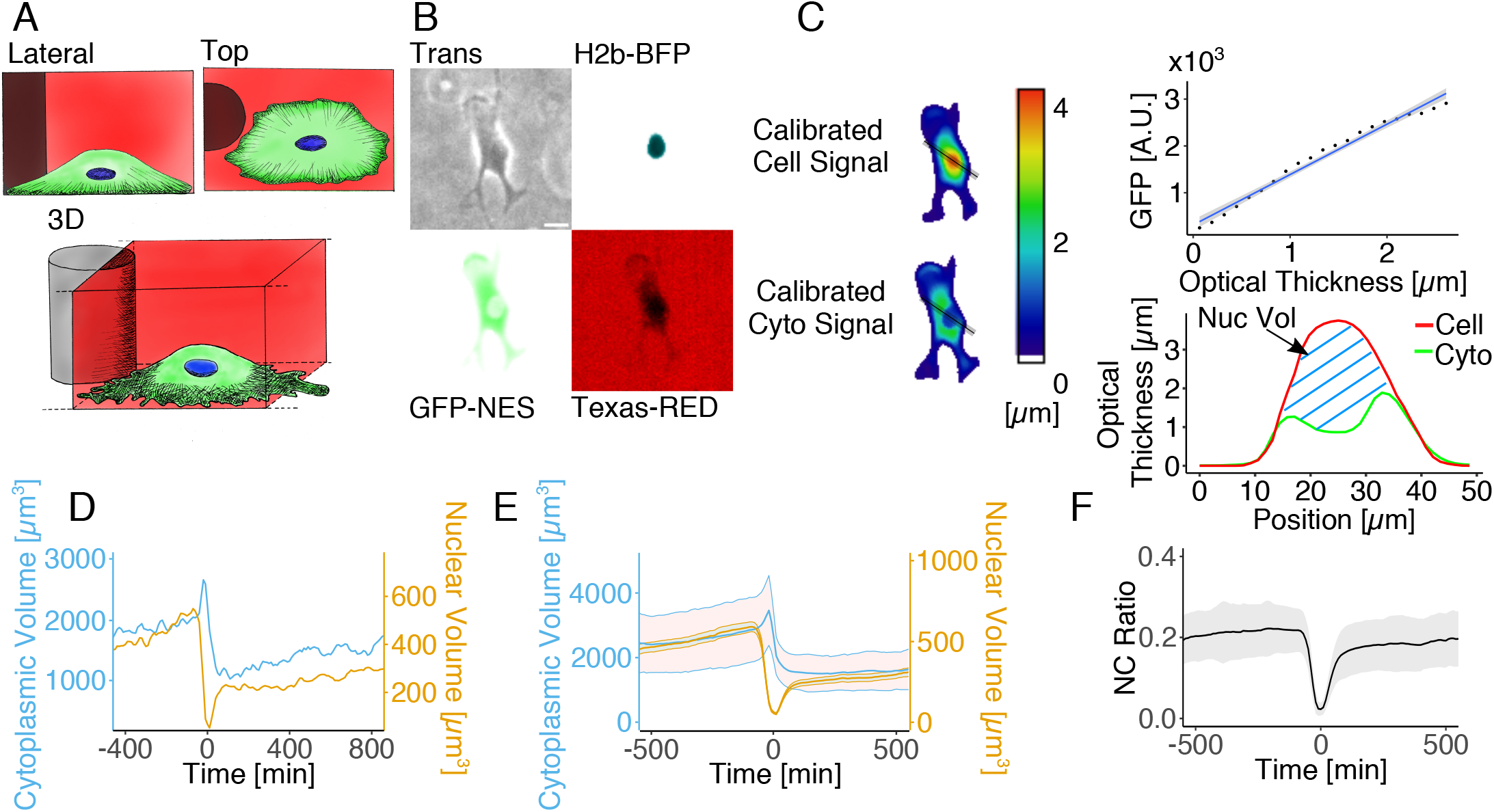
N2FXm development. **(A)** Schematic representation of cell in the acquisition chamber: bottom, 3D rendering; top left lateral view; top right top view. **(B)** Widefield images of a RPE1 cell in the acquisition chamber: transmission light (top left), H2b-BFP (top right), GFP-NES (bottom left) and Texas red (bottom right). Scale bar is 20μm. **(C)** Fluorescence calibration and nuclear volume calculation. On the left: calibrated color-coded images of the cell (top) and cytoplasm (bottom) heights. On the top right: example of experimental calibration curve for the GFP-NES signal. On the bottom right cell and cytoplasm calibrated signal profiles along black line in calibrated images in C. **(D)** Nuclear (blue) and cytoplasmic (yellow) volumes trajectories for a single RPE1 cell. Mean nuclear and cytoplasmic volumes **(E)** and mean NC ratio **(F)** trajectories for RPE1 cell population (n=99). All single curves are aligned to cytokinesis = 0 min.

We employed N2FXm to explore the nucleus-cytoplasm volumetric coupling in nonsynchronized proliferating cells from 5 different epithelial cell lines, normal and transformed: RPE1, MCF10A, MCF7, DCIS.com and MCF10-CA, (Fig. 1D-F, S1L). Surprisingly, cytoplasm and nucleus volume mean temporal trajectories clearly show that NC ratio is not constant over the cell cycle (Fig.1F and Fig. S1L), differently from earlier reports for yeast and from what hypothesized for decades per mammalian cells (*6, 7*).

To gain a deeper understanding of the homeostatic nucleus-cytoplasm coupling, we exploited N2FXm analyzing single-cell dynamic measurements of cytoplasmic and nuclear volumes. Initially, we considered three specific time-points across cell division: I, the instant preceding nuclear envelope breakdown (NEB); II, the onset of cellular roundup at division; and III, the instant successive to the post-mitotic nuclear expansion (PME) (for point identification methods, see “Curve generation and point detection” section in supporting information, Fig. 2A-B and fig S2-A). We found that NEB systematically precedes the onset of cellular roundup by ~10-20 min (Fig. 2C). However, the temporal resolution of our experiments (10 min) was too small to precisely distinguish these two events. Moreover, we found a strong positive correlation between both nuclear and cytoplasmic volumes at NEB and PME, the end of a cell cycle and the start of the following one respectively (Fig. 2D). These results are in line with previous findings and reinforce the idea that “size-memory” mechanisms act to preserve both cytoplasm and nuclear dimensions of the cell population (*12*). Interestingly, while on average cytoplasmic volume almost perfectly halves across cell division (*12*), the nucleus decreases on average by ~2.5 times (Fig. 2E). This implies a non-constancy of the NC volume ratio between NEB and PME (Fig. 2F). These observations also suggested non-trivial homeostatic coupling mechanisms between cytoplasm and nucleus. Then, to test volumetric growth speed of the two cellular compartments, we computed the conditional averages of growth speed (dV/dt) at fixed size and analyzed their average trends. We found that cytoplasm and nucleus grow following different average growth laws. For all analyzed cell lines (confirming previous reports (*12, 14, 15*)) the cytoplasm grows exponentially, with growth speed linearly increasing with cytoplasm size, while the nucleus grows linearly, with constant growth speed at varying nuclear size (Fig. 2G and S2-B). Overall, these results indicate that in mammalian cells the nucleus-cytoplasm volumetric coupling could not be simply defined by a pure osmotic equilibrium, which would lead, instead, to a constant value of the NC ratio (*16, 17*).

**Fig. 2.**
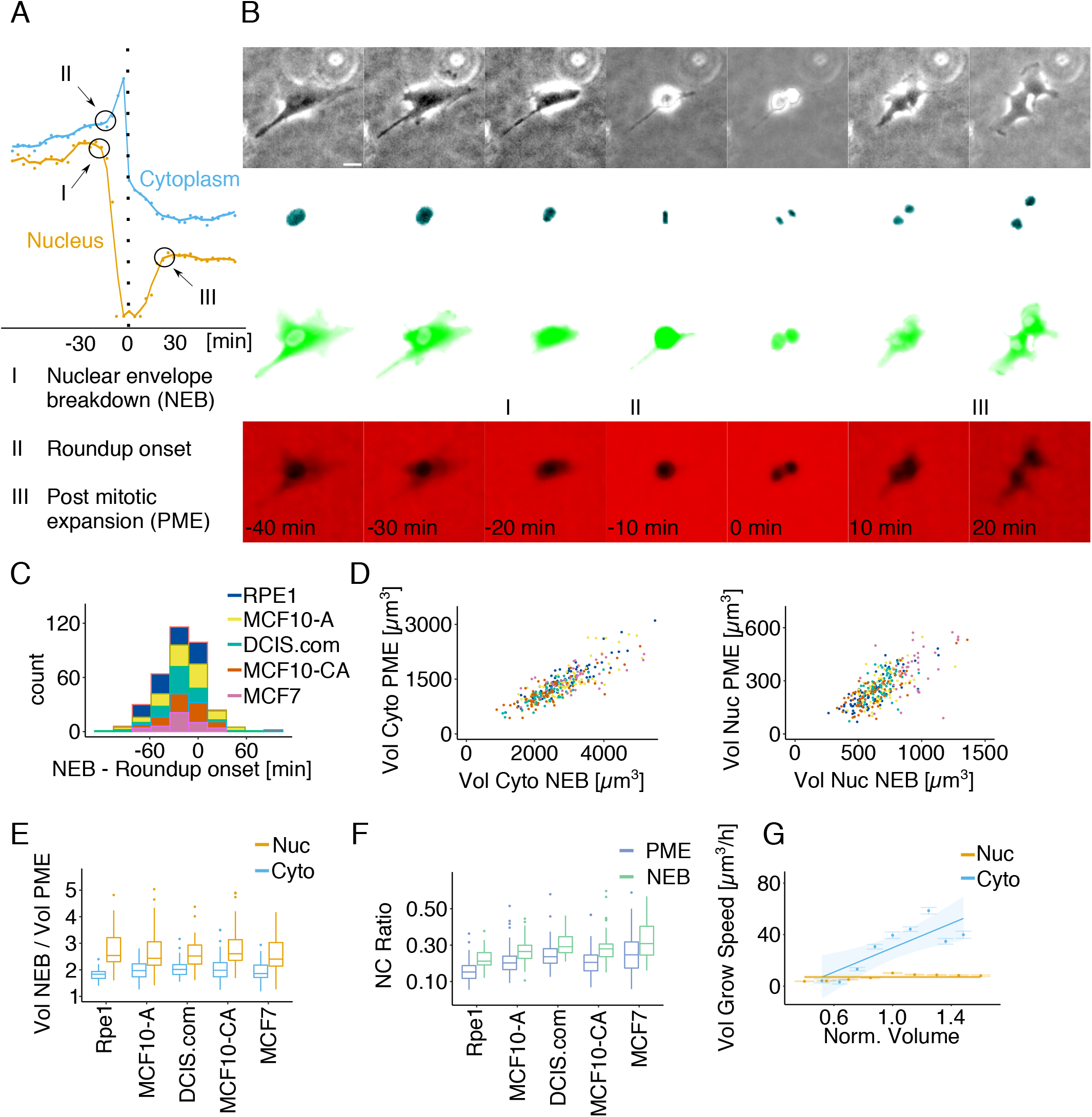
Analysis of nucleus-cytoplasm volumetric coupling across division reveals general relations linking nucleus and cytoplasm volumes. **(A)** Detection example of I (NEB), II (roundup onset) and III (PME). **(B)** Representative images of a RPE1 cell across division. From top to bottom: transmission light, H2b-BFP, GFP-NES and Texas red channels. Scale bar is 20 μm. **(C)** Cumulative distribution of delay between I and II for all the 5 cell lines analyzed. **(D)** Scatter plot linking the cytoplasmic (left) and nuclear (right) volumes at points I and III. **(E)** Boxplot representing the distribution of the volumetric ratio between points I and III for the nucleus and cytoplasm. Mean ±SD. **(F)** Boxplot representing the NC ratio distribution at points I and III. In (C), (D), (E) and (F) RPE1 n=82, MCF10-A n=88, DCIS.com n=66, MCF10-CA n=66, MCF7 n=42. Mean ±SD **(G)** RPE1 cytoplasm and nucleus mean volume growth speed trajectories evaluated in function of the size/volume, n=99; mean ±SE.

To explore the biophysical factors regulating the nucleus-cytoplasm volumetric coupling, we performed a set of experiments on shorter time scales, to osmotically or mechanically perturb RPE1 cells. These stimuli are known to impact on both cell and nuclear volume (*18*–*21*). We coupled these experiments with measurements of the cytoskeletal forces acting on the nuclear envelope (NE), using a FRET sensor based on the LINC complex protein Nesprin 1 (Fig. S3 A-B) (*22*). We also used a set of simple biophysical models to rationalize the results (see SI appendix) (*16, 17*). In the basic version of this model, NC volumes are set only by osmosis and surface tensions are negligible. As expected from a purely osmotic model, hyperosmotic shock induced a substantial shrinkage of both nuclear and cytoplasmic volumes, leaving NC ratio and cytoskeletal forces acting on the nuclear envelope unmodified (Fig. S3A-B). Our model also shows that the same behavior is expected for a non-negligible constant surface tension, with a small correction on the slope, but the expected nuclear volume changes due to external forces are small. Conversely, lowering cytoskeletal forces exerted on NE, as induced in suspended, latrunculin-treated or ROCK inhibitor Y-27632 treated cells (see supporting information), caused a substantial nuclear volume decrease, with only minor effects on the cytoplasmic volume (Fig S3A-B). Clearly, these biophysical changes also lead to significant variations of the NC ratio (Fig. S3A) and falsify the expectations of a purely osmotic model (SI appendix).

Exploiting N2FXm to its full potential, we then analyzed the volumetric dynamic response during three kind of perturbations: hyperosmotic shock, cell detachment and cell spreading. In cells subjected to a hyperosmotic shock, both nuclear and cytoplasmic volumes simultaneously decreased without any detectable lag (at the temporal resolution of our experiments, 2 min), and with negligible effects on the NC ratio (Fig. 3B-D and S3B), once again confirming the osmotic model. Moreover, the force acting on NE, independently assessed, was mostly unaffected along hyperosmotic shocks (Fig. 3E). Instead, during cell detachment, both nuclear volume and force on NE decreased, with no considerable effects on cytoplasmic volume (Fig. 3G-J). Equally, during spreading, nuclear volume and force on NE increased and the cytoplasmic volume decreased (Fig.3L-O) (*23*). Altogether, these results confirmed the osmotic coupling existing between nucleus and cytoplasm, which coherently react to an osmotic perturbation (Fig. 4A). However, they also highlight that external mechanical forces exerted on the NE must be an important regulator of nuclear volume. Indeed, during cell detachment or spreading, or upon cytoskeleton perturbations, nuclear and cytoplasmic volumes are strongly decoupled, with nuclear volume variations coherent with the changes of forces exerted on the NE and not with the changes of cytoplasmic volume (Fig. 4A). Based on the model, we reasoned that this was very unlikely to be a direct effect, as the direction of the effect is opposite to the expectation from cytoskeletal compressive forces (*24*), and the mechanical forces required to change nuclear size would be abnormally large (see SI appendix). Instead, recent findings suggest that NE tension, controlling nuclear pore size, could alter nuclear import rate of small proteins(*25–28*) potentially affecting osmosis (*29*).

**Fig. 3.**
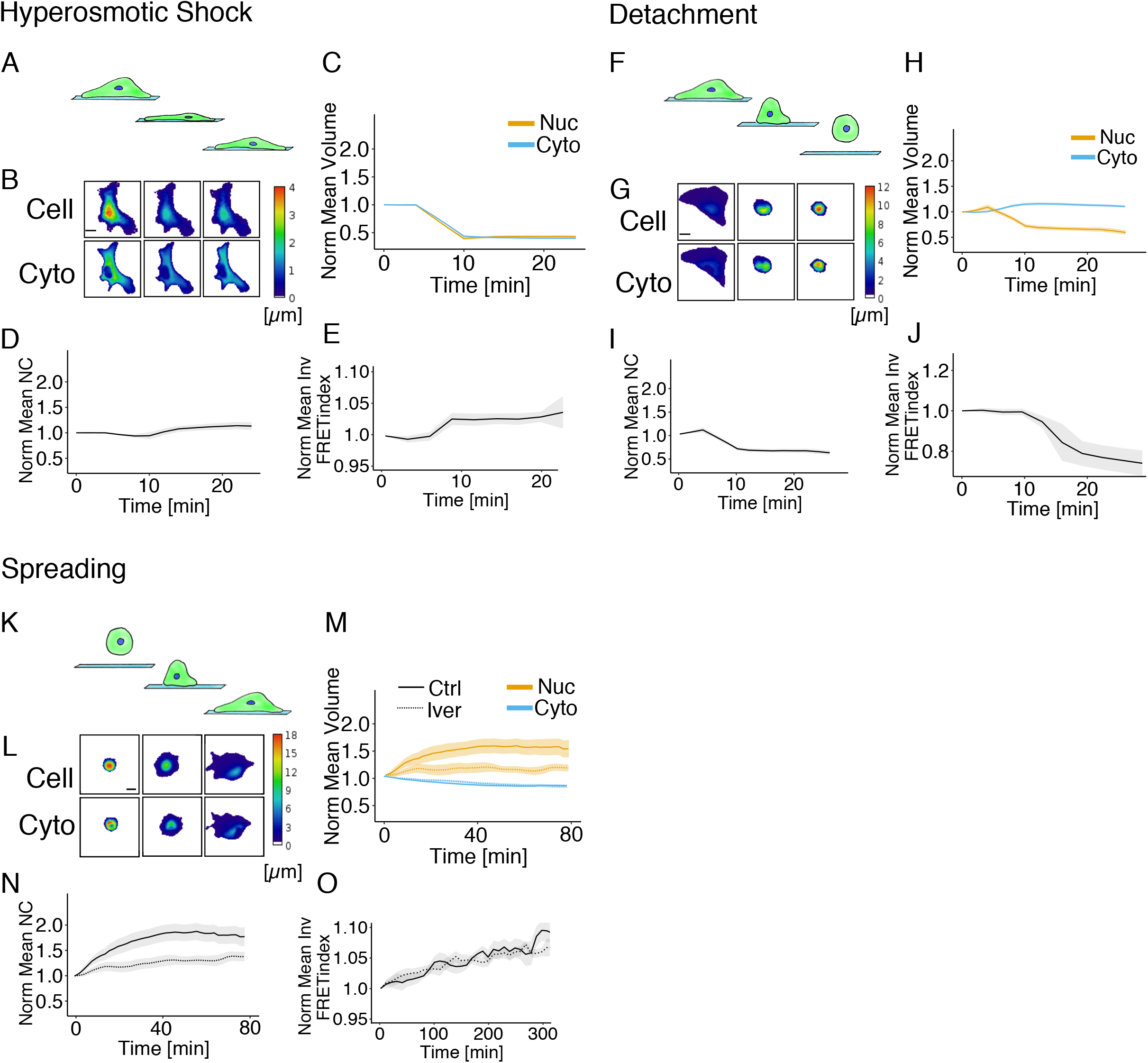
Osmotic, mechanical and active transport contributes in nucleus-cytoplasm volumetric coupling determination. Dynamic perturbations: A-E hyperosmotic shock, F-J detachment and K-O spreading. Representative cell and cytoplasmic height evolution of a RPE1 cell during hyperosmotic shock **(B)**, detachment **(G)** and spreading **(L)** experiments. Scale bar is 20 μm. Mean normalized nuclear and cytoplasmic volume trajectories during hyperosmotic shock (n=24) **(C)**, detachment (n=23) **(H)** and spreading **(M)**. In **(M)**, the dashed line is relative to ivermectin treated cells (n=22), while the continuous line to untreated cells (n=47). Mean normalized NC ratio trajectories during hyperosmotic shock **(D)**, detachment **(I)** and spreading **(N)**. In **(N)**, the dashed line is relative to ivermectin treated cells, while the continuous line to untreated cells. Mean normalized inverted FRET index value during hyperosmotic shock (n=20) **(E)**, detachment (n=10) **(J)** and spreading **(O)**. In **(O)**, the dashed line is relative to ivermectin treated cells (n=10), while the continuous line to untreated cells (n=14).

**Fig. 4.**
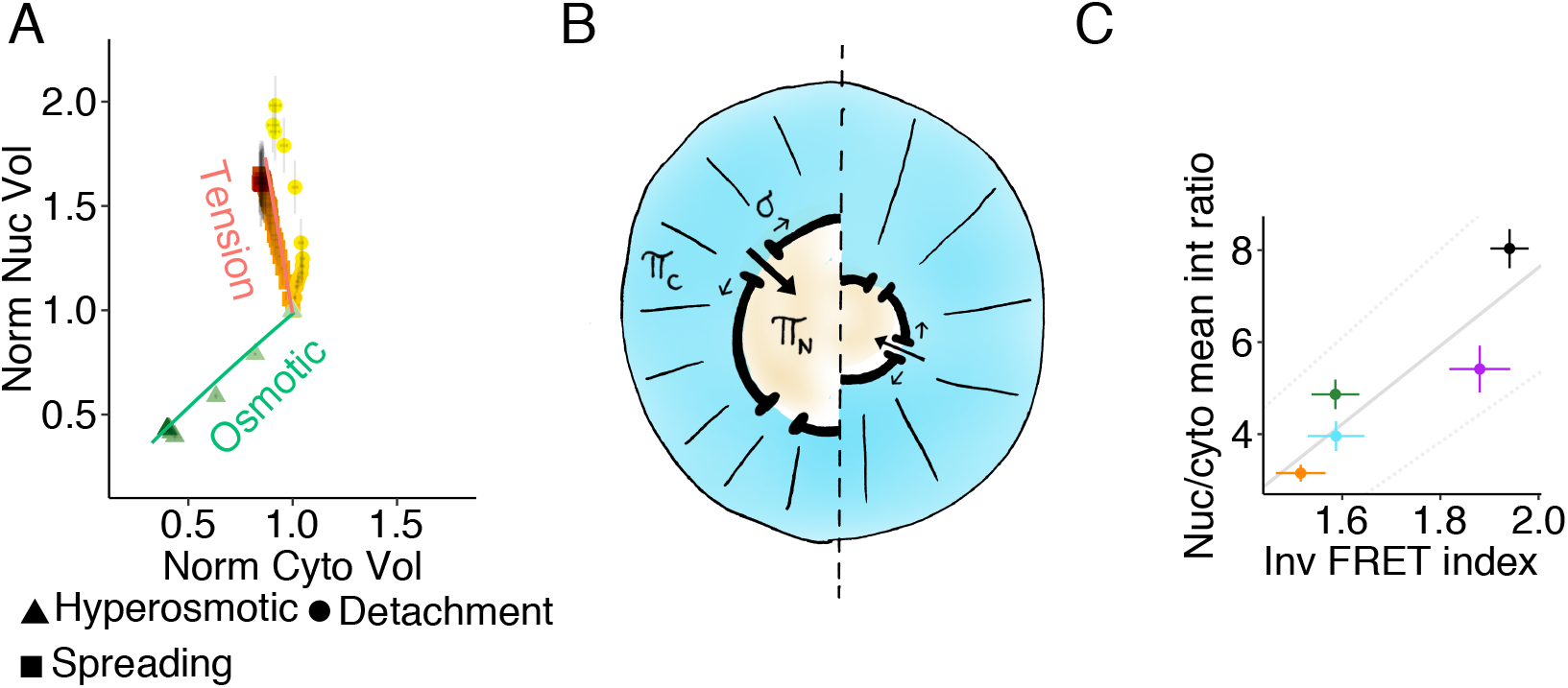
Modeling of the tension-biased regulation of nucleus-cytoplasmic volumetric coupling. **(A)** Scatter plot representing different nucleus-cytoplasm volumetric coupling relationship (variations of nuclear volume as a function of cytoplasmic volume, normalized to resting values) in hyperosmotic shock (green), detachment (yellow) and spreading (orange, untreated cells) experiments (Mean ± SE). Here, time progression is coded with a color gradient (from light to dark). The green and orange solid lines are predictions from the theoretical model with parameter values: 20\% fraction NLS/NES proteins (in absence of applied tension), and an external tension-transport bias of 4 10^3^ m/N (estimated from Andreu et al. 2021). The orange line is a prediction for a variation of the tension caused by external forces σ_ext_ from 0 to 10^−4^ N/m. Reference cytoplasmic volume (for the model) V_C_ = 2000 μm^3^, reference nuclear volume V_N_ = 354 μm^3^.**(B)** Sketch of the theoretical model. Here, π_c_ and π_n_ represent the oncotic pressure of the cytoplasm and the nucleus, respectively, while sigma the NE tension **(C)** Scatter plot of the mean value of nucleus on cytoplasm ratio of the GFP-NLS mean intensity in function of the inverted FRET index (mean ±SE) in the different condition considered.

We explored a model variant where, in addition to the osmotic equilibrium, we added a term reflecting the contributions of the mechano-mediated regulation of nucleo-cytoplasmic transport (Fig. 4B and S3E, and SI appendix). This model, calibrated with data from refs.(*17, 25*), suggests that moderate tension changes may affect nuclear size through this effect (SI appendix and Fig 4A-B and S3E). Hence, we tested this hypothesis experimentally. We first quantified nuclear import efficiency as a function of the force acting on NE, using as proxy the nucleus-cytoplasm fluorescence intensity ratio in cells expressing GFP-NLS protein. We found that perturbations decreasing the extent of forces exerted on NE always decrease nuclear import efficiency (Fig. 4C). We then assessed volume dynamics in spreading cells treated with ivermectin, an inhibitor of the α/β importins transport pathway (*30*), at a concentration affecting nuclear import but unable to perturb none cytoplasmic or nuclear volume, NC ratio or forces acting on the NE (Fig. 4C S3C-D). During spreading, despite as in untreated cells forces on NE increases and cytoplasm volume decreases, importins impairment lead to a significantly slower nuclear volume increase (Fig. 3M-O). Hence, we conclude that the observed force-related volumetric decoupling was mainly linked to a modulation of the nuclear import. This mechanism can tune the nuclear macromolecular content and thereby controls nuclear volume “amplifying” the sensed forces (*29*).

In conclusion, the N2FXm methods, overcoming common limitations on simultaneous dynamic measurements of cell and nuclear volume in living cells, allowed us to decipher fundamental mechanisms of nuclear-cytoplasmic volume coupling. Unexpectedly, we found that in mammalian cells, cytoplasm and nucleus grow differently, exponentially *vs* linearly. This observation already implies that simple osmotic coupling alone cannot explain nuclear volume dynamics, and more complex homeostatic process must be in place. Our results also show that one of them is related to a sophisticated mechano-sensing mechanism biasing nuclear transport. Indeed, forces exerted by the cytoskeleton, affecting NE tension, impact on nuclear pores size (*26–28*). This, in turn, regulates nuclear-cytoplasmic transport of NLS tagged proteins, affecting osmosis. Accordingly, specific transcription factor such as YAP, key regulator of organ growth and regeneration as well as of mechanotransduction, also are affected by this mechanism, establishing intriguing links between biophysical and regulatory pathways to be explored in the future. Finally, nuclear volume regulation, controlling concentration of nuclear components and translocation of transcription factors, impacts chromatin state and nuclear organization, broadly affecting cell activity.

## Acknowledgments

The authors would like to thank IFOM Imaging Facility for help with performing experiments and the clean room facilities of the Binning and Rohrer Nanotechnology Center (BRNC) for contributing in mold fabrication. We would like to acknowledge Clotilde Cadart, Alessandro Marcello, Kristina Havas Cavalletti, Nils Gauthier and Giorgio Scita for helpful comments and discussions and thank Gilda Nappo for illustrations.

## Author contributions

FAP was responsible of the project. FAP, AP and EP designed, carried out and interpreted experiments. AP developed the miniNesprin1 FRET sensor. FMP, under the supervision of AF, participated to the design and produced the microfluidic measuring chamber. SL, under the supervision of IR, performed cellular biology experiments. MC and AP designed and cloned the plasmids. DV provided DAAM particles and helped designing experiments. MP and MCL designed and interpreted experiments. MCL and OMR developed the mathematical model. FAP carried out images, data and statistical analysis and wrote the paper with PM and MCL. PM supervised the study.

## Data and materials availability

All data are available in the main text or the supplementary materials.

**Supplementary Materials**

Materials and Methods

Supplementary Text

Figs. S1 to S3

References (*##-##*)

Movies S1

Appendix

### Supplementary Materials for

#### Cell Culture

All culture media were supplemented with 1% L-glutamine and 1% penicillin-streptomycin. hTert RPE1 ATCC cells were cultured in DMEM-F12 supplemented with 10% FBS. MCF10-A ATCC and DCIS.com cells (Barts Cancer Institute, Queen Mary University of 924 London, UK) were cultured in DMEM-F12 supplemented with 5% horse serum, 10μg/ml human insulin, 0.5mg/ml hydrocortisone and 20ng/ml epidermal growth factor (EGF). MCF10-A medium were cultured using the same DCIS.com medium and further supplemented with 100 ng/ml cholera toxin. MCF10-CA cells (Barts Cancer Institute, Queen Mary University of 924 London, UK) were cultured in DMEM-F12 supplemented with 5% horse serum, 1.05 mM calcium chloride and 10mM HEPES. MCF7 NCI cells were cultured in RPMI 1640 medium. All the cell lines were grown at 37 °C and in humified atmosphere with 5% CO2.

#### Cell transfection, lentiviral production and cell transduction

Lentiviral particles were produced as described here (31). Briefly, 3×10^6^ HEK293T cells were transfected with pCDH lentiviral plasmids encoding H2B-BFP or 2X_NES-GFP (nuclear exportation signal), or MiniNesprin1_FRET sensor, together with psPax2 and pMD2.G vectors with a ratio of 4μg: 2μg: 1μg using Polyethylenimine (PEI) as transfection reagent. 24h later medium was removed, and virus collection performed after 24/48/72h. Viral aliquots were pulled together and filtered with 0.45μm filters. 2×10^5^ cells were transduced with 1ml of fresh virus supplemented with Polybrene (8(×)g/ml) through 20’ of centrifugation at 2000rpm, then seeded in 6 well plates.

Cell transfection of hTERT_RPE1 cells was performed using Neon Transfection System (Thermo Fischer Scientific) following the manufacturer’s protocol. Expression of nuclear GFP was obtained through a plasmid encoding a NLS (nuclear localization signal) tagged eGFP (Addgene #67652).

#### N2FXm Chip fabrication

The N2FXm chip is a microfluidic device composed by two “acquisition” chambers connected by “loading” channels (Fig. S1A). Acquisition chambers are 18 μm in height and used to image cells for volume measurements. Reservoir channels are higher (~150 μm) and ensure a continuous nutrient delivery to cells in the acquisition chambers (Fig. S1A). Microstructured silicon mold were used to create the N2FXm chip. The template wafers were fabricated at the cleanroom facilities of the Binning and Rohrer Nanotechnology Center (BRNC) using a standard two step-photolithography process (*32*). First, the pillars in the central chamber were etched until a final depth of 18 μm. Finally, the lateral channels were added by crosslinking a 150 μm thick layer of SU-8 resist. The resulting silicon wafers were passivated as follow: soaking for 30 minutes in a silanization solution I (Sigma-Aldrich, USA), then 10 minutes in hexane (Sigma-Aldrich, USA), 10 minutes in 1-octanol (Sigma-Aldrich, USA), and finally rinsed with acetone and deionized water. (Silicon master fabrication). The microfluidic chip was then fabricated in PDMS (Sylgard 184) through standard replica molding. Briefly, the PDMS precursor was mixed with the crosslinker (10:1) and poured on the silicon mold, degassed for 1h in a vacuum bell and then cured for 3h at 90 °C. Once demolded, 1.5 mm diameter punches were made in correspondence of the inlet and outlet of the chip. The chip was then treated for 1 min with oxygen plasma and irreversibly bonded to a 35 mm bottom-glass petri dish (Mattek). To reinforce the bonding treatment, the bonded device was heated for 30 min at 40 °C. To prevent medium leakages, two PDMS cubes (height ~3 mm) were punched (2 mm diameter) and bonded in correspondence of the inlet and outlet. The day before the experiment, the chip was retreated for 1 min with oxygen plasma, filled with fibronectin at the appropriate concentration (10 μg/ml for RPE1 cells, 20 μg/ml for all the other cell lines) and incubated at RT for 1h. The chip was then rinsed with PBS, filled with the appropriate medium and incubated overnight at 37 °C. The day of the experiment, cells were injected in the chip at ~300×10^3^ cells/ml. The petri dish was humified (filled with medium) to prevent medium evaporation from the chambers during the experiment. For all the experiments excepting spreading, after 3-6 h from cell injection (the time necessary to ensure a good adhesion of a specific cell line), the culture medium was replaced with a medium containing 70kDa Texas Red Dextran at 1mg/ml (Thermofisher Scientific) and, after additional 3h, the microfluidic system was imaged at the microscope. In the case of spreading experiments, cells were instead directly resuspended in the Texas Red Dextran containing medium, injected in the chip and imaged ~15 min later.

#### N2FXm for Nuclear Volume measurement

Cell and nuclear volumes were measured imaging GFP-nes and H2B-BFP positive cells in the N2FXm chip filled with a fluorescent dye (Dextran Texas red 70 kDa Neutral, Invitrogen) not internalized by cells (see “Chip fabrication” section, Fig. S1-A). We acquired images in 4 channels: transmitted light, red (dextran), green (GFP-NES) and blue (nuclear staining). Images were analyzed using ImageJ. First, dextran field illumination homogeneity was corrected dividing the original image by a correction/normalization image, which was obtained by dividing the smoothed original image (or the maximal projection or multiple frames, in case of acquisitions longer than 1 time point) for its mode. Here, the smoothing was obtained applying a median filter with a radius value ranging from 35-50, which corresponded to the minimum value needed to eliminate the contribute of the pillars (i.e. that appeared as black circles in the image) from the mode calculation of the dextran channel. Then, as in Zlotek-Zlotkiewicz et al., drop of dextran intensity in correspondence of cells was used to calculate cells volume (*10*). Any object in the chamber, occupying a specific space, excludes the red fluorescent dextran from this volume. Dextran fluorescence intensity can be scaled between two points of known height: zero, where the chamber is empty and the maximum, the known chamber height (18um), in correspondence of the pillars sustaining the chamber roof (as schematically showed in Fig. S1B). This calibration allowed defining the optical thickness of cells in the chamber at each pixel and was performed independently for all fields of view and all time points. Although optical thickness at each pixel can differ from cell height, its integration over a slightly enlarged cell area, here automatically defined from the GFP-NES fluorescence signal, well quantify the cell volume (Fig. S1C-E) (*10*).

For measuring nuclear volumes, a second calibration was introduced. Cell nuclei were negatively stained with the ectopic expression of a green fluorescent protein coupled with a nuclear export signal (GFP-NES), which localizes in the entire cytoplasm except the nucleus, also marked with H2B-BFP (see “Cell transfection, lentiviral production and cell transduction” section). Since we assume that the intensity of the GFP signal is linearly proportional to the volume of the corresponding portion of cytoplasm, it can as well be calibrated according to the one of calibrated dextran. The GFP calibration was then performed fitting the conditional average of GFP intensity in the cytoplasm (all cell excluding the nucleus) at fixed optical thickness (Fig. S1C-E). Cytoplasm and nucleus were automatically defined by combining the GFP and BFP signals (Fig. S1E). Linear fitting was carried out in R using the “robustbase” package [Maechler M, Rousseeuw P, Croux C, Todorov V, Ruckstuhl A, Salibian-Barrera M, Verbeke T, Koller M, Conceicao EL, Anna di Palma M (2021). *robustbase: Basic Robust Statistics*. R package version 0.93-9, http://robustbase.r-forge.r-project.org/]. Importantly, to correct for possible signal fluctuation or variation of GFP expression, GFP calibration was performed at each timepoint and for each analyzed cell. In order to strengthen the calibration, we introduced also the possibility to perform and mediate it over a variable number of timepoints (nzz). Volume trajectories relative to asynchronous cells along the cell cycle (Fig. 1D-F and Fig. S1M) were obtained setting nzz=3 (see supporting video S1). Spreading, detachment and hyperosmotic shock dynamic experiments were analyzed with nzz=1.

Finally, to measure nuclear volume, the calibrated GFP signal was integrated within the nuclear region, and the resulting value subtracted from cell volume evaluated over the same domain (see materials and methods and Fig. S1-E). Cytoplasmic volume was calculated by subtracting the nuclear volume from the entire cell volume and the NC ratio as the ratio between nuclear and cytoplasmic volumes.

#### Live imaging experiments

All live imaging experiments were performed at 37 °C and 5% CO_2_ atmosphere. FXm experiments were performed using a LEICA widefield DMI8 inverted system equipped with a HC PL Fluotar 10x NA=0.32 (Leica, #506522) objective. The excitation source was a LED illumination. Images were acquired with a sCMOS Andor Neo 5.5 camera.

FRET experiments and 3D reconstruction were both performed with an inverted LEICA confocal SP8 equipped with a HC PL APO 40×, NA=1,30 OIL immersion (Leica, #506358) or a HC PL APO 63×, NA=1,40 OIL immersion (Leica, #506350) objective, respectively.

GFP-NLS nuclear-cytoplasmic localization was obtained with an inverted spinning disk (CSU-X1 Perkin Elmer Ultraview) Nikon Ellipse Ti, equipped with Nikon PLAN Fluor 40X 1.3 OIL objective and a Hamamatsu EM-CCD C9100 camera.

#### DAAM particle measurement and N2FXm precision

Poly-acrylamide-co-acrylic acid particles (DAAM particles) were used for evaluating the precision of our technique. In particular, if internalized by the cell, DAAM particles are excluded by the cytoplasmic staining (while increasing the total cell volume) and their volume could be then measured using our technique. DAAM particles were preferred to glass beads because the refractive index (RI) of the latter (RI >1.47) was too different from that of the water (RI~1.33) and generated optical artifacts, while DAAM RI ranged between 1.33 and 1.34 (*11*). DAAM particles were produced as previoulsy decribed with a mean diameter of 3.5 μm (*11*). DAAM particles were resuspended at 50×10^6^ beads/ml in the RPE1 cells medium and injected in the acquisition chamber containing adhered RPE1 cells. After 4 h from particle injection, culture medium was replaced with Texas red dextran (1mg/ml) containing medium. To estimate the error, volumes of the internalized particles were quantified with our technque (e.g. measured volume) and compared with the ones obtained through a geometrical calculation (e.g. expected volume) (Fig. S1G-I). Here, because of the particle spherical shape, expected volumes could be calculated measuring particle diameter (averaged between two measurements). This procedure was performed manually and not by an automatic measurement.

Moreover, we compared nuclear volume distribution measured with the N2FXm with the one obtained by means of confocal 3D reconstruction. 3D reconstruction was performed by imaging RPE1 cells in the same culture condition used for the N2FXm experiments (see “chip microfabrication” section). Cells were then acquired along z with a step size 0.3 μm. The stacks were then analyzed with the 3D object counter plugin of ImageJ. From this analysis, we did not find statistical differences between the volumes obtained with the two methods (Fig. S1F).

#### Drug and adhesion experiments

All the drug experiments were performed for both volume (N2FXm) and nuclear tension (FRET) measurements. For N2FXm, drugs were added at the proper concentration to the specific medium (containing the fluorescent dextran) and injected in the device containing adherent cells (as described in the “Chip fabrication” section). In GFP-NLS localization experiments, all the condition were explored excepting for hyperosmotic shock.

For perturbing cytoskeleton and cell mechanics, latranculin-A (LatA, Sigma Aldrich) and Y-27632 (Y27, Sigma Aldrich) were added to the culture medium at 1μM and 500μM, respectively. Cells were then imaged after 1h (for LatA) or 12h (for Y27). For hyper-osmotic shock experiments, D-sucrose (Carlo Erba reagents, Dasti Group) was resuspended in pure water at 500 mM and, to perform dynamic measurements, added to cells during imaging. Detachment experiments were performed by adding EDTA to a serum-free culture medium at (0.05%). As in the case of osmotic shock experiments, EDTA containing medium was added during imaging for obtaining dynamic measurement. Spreading experiments were conducted by imaging cells ~15 min after seeding. Here, untreated (control) cells and cells pre-treated with ivermectin (Sigma Aldrich, 10 μM for 24 h) were used.

In N2FXm dynamic experiments (i.e. hyper-osmotic shock, detachment and spreading), cells were acquired each 2 min (see supporting video S2). In FRET dynamic measurements cells were acquired each 3 min during osmotic shock and detachment, while during spreading each 10 min. The lower acquisition rates used for spreading FRET experiments were justified by the need to perform z-stack acquisitions (z step size=1μm) for accounting cell movements along the z axis.

#### Curve generation and point detection

All the figures and statistical analysis were performed in R. Packages used were: “robustbase”, “ggplot2”, “gridExtra”, “rcarbon”, “ggsignif”, “rcarbon”, “rsvg”.

All the plotted dynamic datasets were smoothed using the “runMean” function (“fill” smoothing option on 3 points). In all the smoothed datasets, eventual NA were removed using the “na_interpolation” (linear option) function.

To generate mean volume and ratio trajectories of non-synchronized growing cells (Fig.1E-F and S1M), single curves were first temporally aligned and then mediated over time. Here, cytokinesis was used as reference time point (represented as the time “0 min”), which was manually identified for each cell analyzed from the acquired images.

Nuclear envelope breakdown (NEB), postmitotic expansion (PME) and roundup onset points relative to single cell trajectories were automatically detected (Fig. 2A and S2A). The detection was performed as follows:

NEB: This point was detected by combining the first and the second differential of smoothed nuclear volume data (runMean function, “fill” smoothing option on 3 points). In particular, the NEB was defined 1 point before the last positive value of the first differential, which was considered in a 10 points wide window preceding the maximum value of the second differential before division. The second differential was analyzed in a 15 points window before division.
PME: 5 points successive to the minimum value of the second differential of smoothed nuclear volume data (runMean function, “fill” smoothing option on 15 points or on the maximum length of the nuclear trajectory after division, if the latter was lower than 15), considering a 15 points wide window after division.
Roundup onset: 2 points before the maximum value of the second differential of the smoothed cytoplasmic volume data (runMean function, “fill” smoothing option on 15 points or on the maximum length of the nuclear trajectory before division, if the latter was lower than 15), considering a 15 points wide window before division.

All the detected points were manually checked.

To generate the normalized dynamic mean curves relative to volumes, NC ratio and inverted FRET index (fig. 3C-E, H-J, M-O) during spreading, detachment and osmotic shock experiments, single curves were first smoothed by means of the runMean function (“linear” smoothing option on 3 points). Successively, data relative to each curve were divided by their first value to be normalized. Finally, all the curves were mediated over time.

#### Growth Speed analysis

Cytoplasm and nuclear growth modes were obtained by analyzing the volume growth speed (dV/dt) in function of the volume (V). Here, because of the already reported high variability of single cell trajectories (*13*), growth modes were evaluated considering the population averages, and in particular the trend of the conditional averages. In this analysis, the two limit cases are represented by the linear and the exponential growth modes, characterized by a constant or linearly increasing value of the growth speed in function of the volume, respectively.

First, single cell volumetric data (i.e. nucleus and cytoplasm volumes used for obtaining the mean volumes trajectories of Fig.1E and Fig. S1L) were smoothed with a central average algorithm considering a sliding window of 20 frames. To calculate the growth speed (dV/dt), a linear fit of single-cell volume curves as a function of time were performed on sliding windows of 12 frames. To get the average population behavior, growth speeds were mediated within 9 volumetric intervals defined to cover the entire volumetric dynamic range. To plot together cytoplasmic and nuclear volume growth speeds curves, volumes were normalized respect to their mean values.

#### Nuclear Envelope tension analysis and synthesis of a novel cpst-FRET sensor

The extent of cytoplasmic forces acting on the Nuclear Envelope (NE) was measured exploiting a novel circularly permutated stFRET (cpstFRET) (*28*) sensor. The sensor has been built using Nesprin 1 protein as backbone and named Mini-Nesprin 1. A detailed explanation about how this construct was created is reported here ((*20*)). Briefly, N-terminus Nesprin 1 (1521bp, aa 1-507) and C-terminus (1422bp, aa 8325-8797) coding sequences containing Nesprin 1 Calponin (CH-CH) and KASH domains respectively were synthesized exploiting IDT gBlocks Gene Fragments Technology. A circularly permutated stFRET (cpstFRET) probe was inserted between the two blocks. Final coding sequence of Mini Nesprin1 cpstFRET sensor was obtained through molecular cloning and inserted into a pCDH lentiviral plasmid. Image acquisition was performed using a Leica TCS SP8 confocal microscope with microscope with Argon light laser as excitation source tuned at 458 nm and HC PL APO CS2 ×40/1.30 oil-immersion objective.

Emission signals were captured at 490-510nm and 515-535nm, and inverted FRET index calculated as their ratio. Data were analyzed using in-house ImageJ macro.

#### GFP-NLS localization analysis

Nucleoplasmic transport was evaluated by quantifying the nucleus on cytoplasm mean intensity ratio of GFP-NLS protein. All the images were analyzed with ImageJ. Here, higher ratios indicate higher active molecular nuclear import.

---

**Fig. S1.**
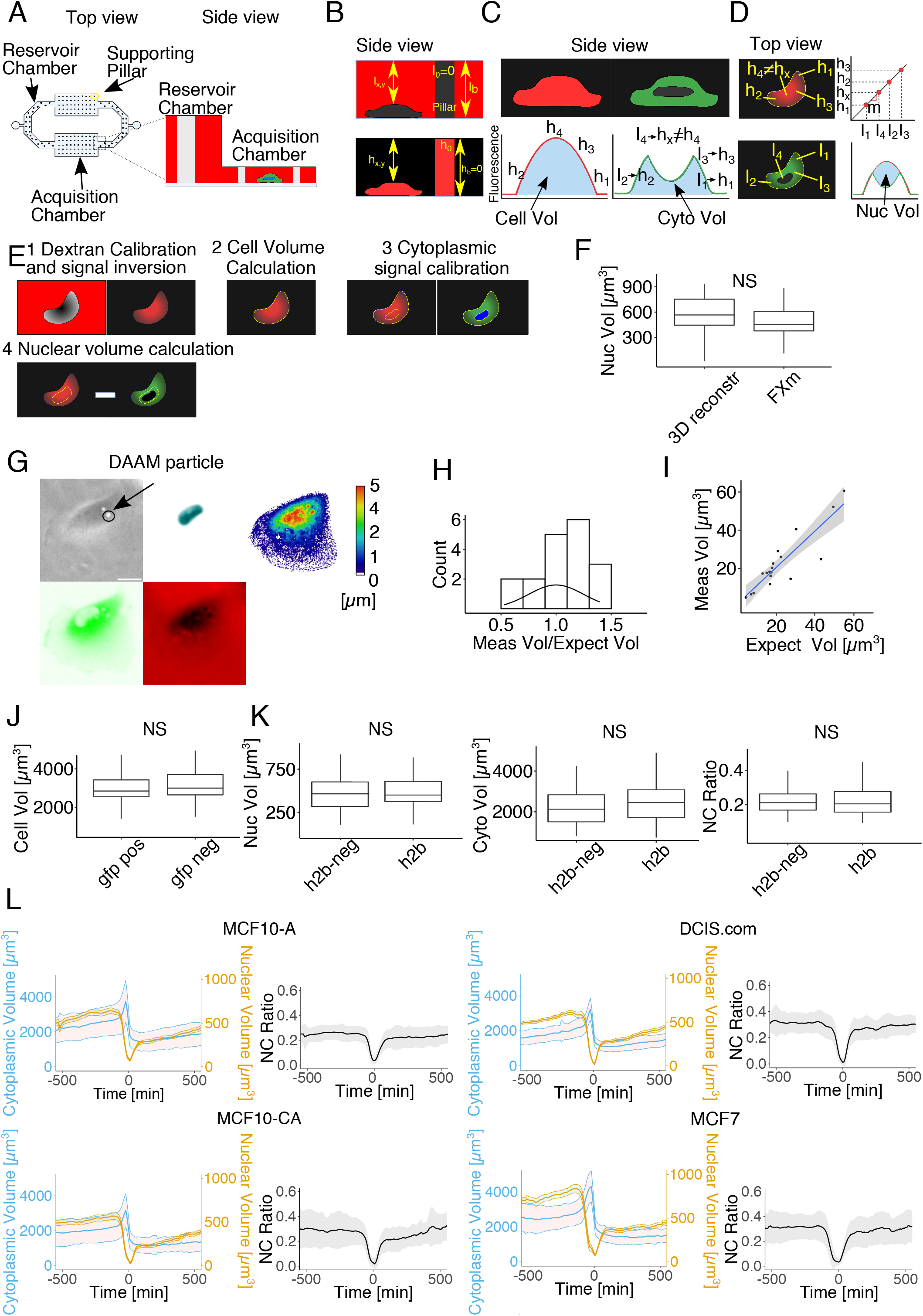
Technique development and mean volume curves generation. **(A)** Sketch of the microfluidic device for FXm acquisition (top and side view). In the side view, the connection between the reservoir chamber and the acquisition chamber is showed. Schematic representation of: **(B)** the first calibration route for cell volume measurement, **(C)** the calibrated cell height profile (left) and the non-calibrated cytoplasmic height profile (right), **(D)** the second calibration route and nuclear volume calculation (bottom right). **(E)** Sketch of the image analysis workflow: (1) first calibration is performed, (2) by defining cell edges from the cytoplasmic staining, cell volume is calculated, (3) combining nuclear and cytoplasmic fluorescent signals, the cytoplasmic region between cell edges and the nuclear region is automatically defined and the second calibration is then performed, (4) nuclear volume is computed by subtraction considering both cell and cytoplasmic calibrated signals in a region equal or bigger than the nuclear region (user defined). **(F)** Nuclear volume distributions comparison considering volumes measured with the FXm (n=95) or with confocal 3D reconstruction (n=50). Welch Two Sample t-test gave p=0.30. **(G)** 10x image of a cell internalizing DAAM particles. Here, transmission, BFP, GFP, Texas red and the associated calibrated image (on the right) of a cell are showed. Scale bar 20 μm. **(H)** Distribution of measured on expected values ratio (n=18). Measured on expected value ratio distribution was compared with a normal distribution centered on 1 with a one sample t-test, giving a p value=0.39 and indicating that the error relative to our measurement was not systematic. **(I)** Scatter plot of DAAM particle “measured vs expected” volumes. Linear fit coefficients: slope=0.94, Pearson correlation coefficient=0.89, R^2^=0.78. To assess if the introduction of exogenous fluorescent proteins (i.e. GFP-NES and H2B-BFP co-infection) altered cellular dimensions, we compared cell volume of differently stained cells. To evaluate eventual effects of the GFP-NES, we compared cell volumes of stained (n=86) and non-stained (n=42) populations. Both cell-types were analyzed in the same device and during the same acquisition. Cell volume was evaluated for all the tested cells 480 min before cytokinesis. Welch Two Sample t-test gave p=0.39 **(J)**. For what concern nuclear staining, instead, we compared nuclear, cytoplasmic volumes and the relative nc ratio distributions of double stained cells (2NES-GFP and H2B-BFP, our control, n=95) with cells presenting only the cytoplasmic staining (H2B-neg, n=91). We did not found significant differences (p=0.089 and p=0.32, p=0.11, respectively) **(K)**. **(L)** Mean volume and mean NC ratio trajectories across division of MCF10-A (n=109), DCIS.com (n=78), MCF10-CA (n=82) and MCF7 (n=62) cells. Time frame= 10 min. Mean ± SD.

**Fig. S2.**
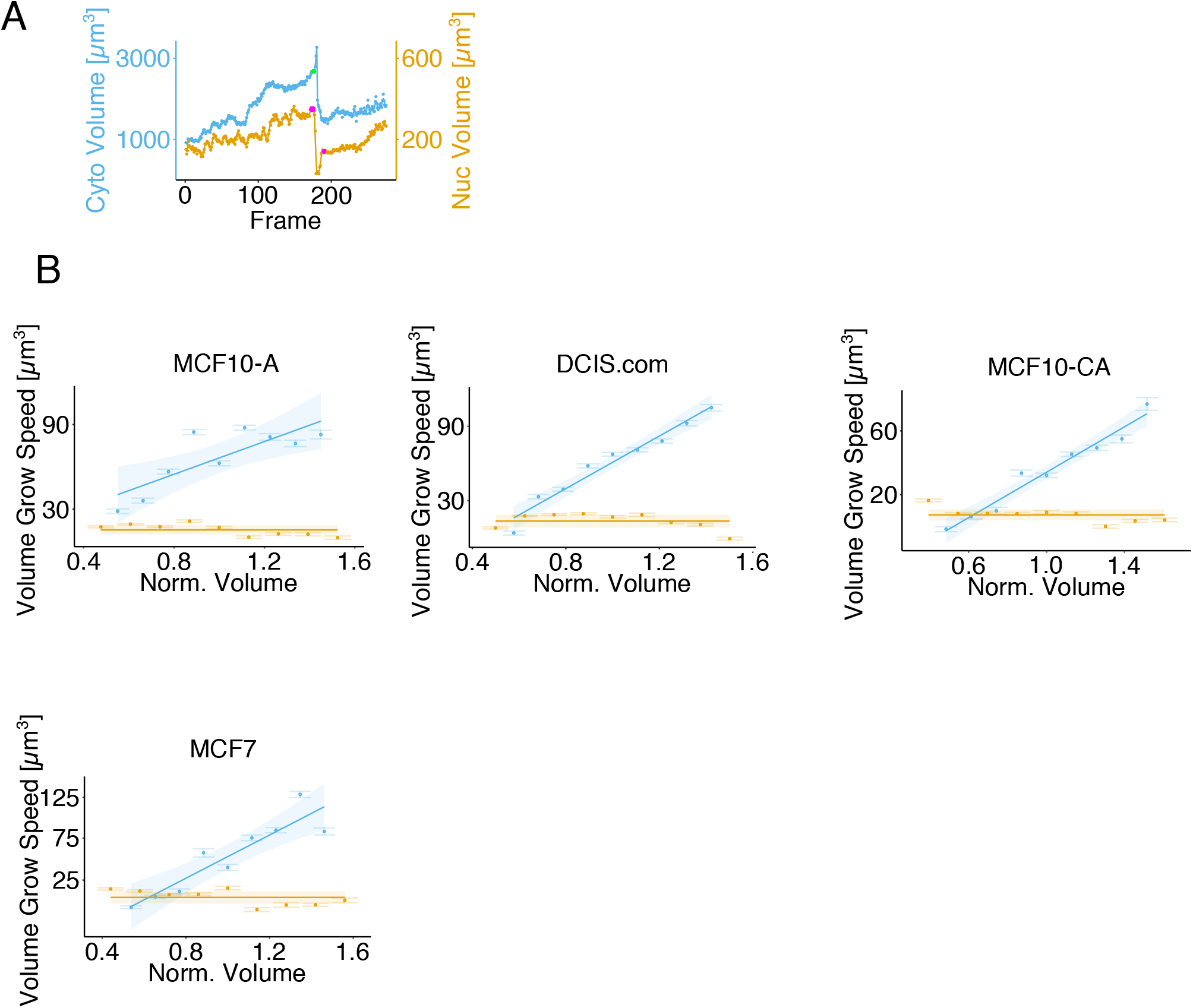
Dynamic analysis and growth curves generation. **(A)** Point detection example on a single cell trajectory. Here, points in magenta on the nuclear volume curve represent NEB (before division) and PME (after division), respectively, while green point on the cytoplasmic curve represents the roundup onset. **(B)** Mean growth speed curves of the nuclear and cytoplasmic volumes (in yellow and blue, respectively) in function of their size. Here, for MCF10-A n=109, for DCIS.com n=78, for MCF10-CA n=82 and for MCF7 n=62. Mean ± SE

**Fig. S3.**
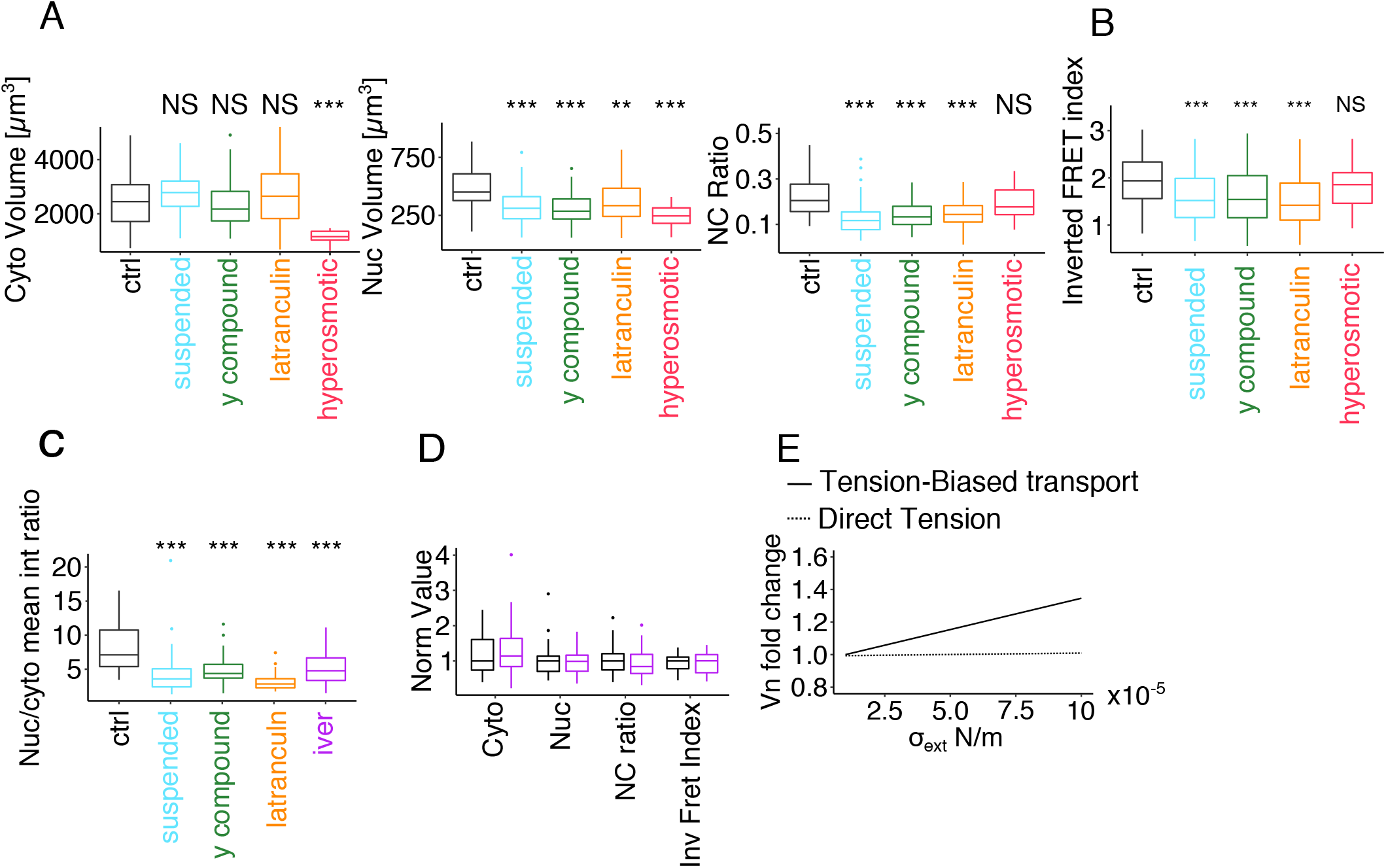
Mechanical nuclear volume regulation. **(A)** Nuclear and cytoplasmic volumes distribution for untreated adherent cells (ctrl, n= 95), untreated detached cells (n= 77, p_nuc_= 0.00014, p_cyto_= 0.071, p_ratio_= 2.35^−7^), Y27 treated cells (n= 64, p_nuc_= 1.54 ^−6^, p_cyto_= 0.89, p_ratio_= 3.90 ^−10^), latranculin treated cells (n= 85, p_nuc_= 0.0070, p_cyto_ 0.16, p_ratio_= 2.55^−5^ and for cells subjected to hyperosmotic shocks (n=24, p_nuc_= 2.27^−13^, p_cyto_= 2.41 ^−9^, p_ratio_= 0.070). **(B)** Inverted FRET index (i.e. indirect measurement of nuclear envelope tension) distribution for untreated adherent cells (ctrl, n= 156), untreated suspended cells (n= 91, p=9.66^−7^), Y27 treated cells (n= 119, p= 3.57^−8^), latranculin treated cells (n=129, p= 1.58^−10^) and for cells subjected to hyperosmotic shocks (n= 92, p= 0.051). Here, Y27 compound and latranculin were used to assess the effects related of stress fibers and contractility, respectively. **(C)** Distribution of the nucleus on cytoplasm ratio of the GFP mean intensity for untreated adherent cells (n=58), untreated detached cells (n=38, p=2.07^−11^), Y27 treated cells (n=44, p=4.52^−08^), latranculin treated cells (n=45,p=2.2^−16^) and ivermectin treated cells (n=44, p=0.00016 **(D)** Normalized volumes (nuclear and cytoplasmic), nc ratios and Inverted FRET index distributions relative to untreated (black) and ivermectin (magenta) treated cells. Data were normalized for the median/mean of the considered variable (i.e. nuclear volume, cytoplasmic volume, nc ratio, inverted FREt index), which was calculated over the control dataset. Here, n_ctrl_ = 45, n_iver_ = 45. p_nuc_= 0.19, p_cyto_=0.83, p_ratio_= 0.16 for volumes data. For Inverted FRET index distribution n_ctrl_= 56, n_iver_= 85, p=0.95. Statistical analysis was performed by means of the welch two sample t-test function in R. NS for p>0.05, * for p<0.05, **for p<0.01, *** for p<0.001. **(E)** Quantitative prediction of the developed theoretical model describing the effects on nuclear volume mediated by a transport biased mechanism (continuous line) or by a direct force (dashed line). The plot reports the predicted fold change in nuclear volume (y axis) for increasing applied extensile tension *σ_ext_*. Parameter values: cytoplasmic volume V_C_ = 2000 *μ*m^3^, 20% fraction NLS/NES proteins in absence of applied tension, and a tension-transport bias of *η* = 4 10^3^ m/N (estimated from Andreu et al. 2021 (*26*), see SI appendix). This model neglects direct effect of extensile tension on nuclear size. The dashed line instead refers to the direct effects of applied extensile tension, which lead only to a mild increase of nuclear volume. The plot reports the predicted fold change in nuclear volume (y axis) at fixed cytoplasmic osmotic pressure for increasing applied extensile tension *σ_ext_*. In the plot, nuclear volume increases only by 0.1%. Parameter values: *α* = 2/3 (fraction of osmotically active proteins that are in the cytoplasm), basal nuclear surface tension *σ*_0_ = 10^−6^ N/m, cytoplasmic osmotic pressure *⊓_C_* = 4 kPa. See SI appendix for a detailed discussion of the models and the parameters.

**Movie S1.**

Example of cell and nuclear volume curves generation for a RPE1 cell dividing twice during the acquisition. Here, the 3 fluorescent channels are merged (BFP, GFP and Texas Red) and the all the ROIs defining the cell area reported.

## SI appendix for Pennacchio et al. Osmo-mechanical equilibrium model for nuclear volume

This SI appendix describes the mathematical model used to complement our understanding of our experimental data.

### Model ingredients

The key ingredient of the model is the assumption that osmotic pressure plays a primary role in setting nuclear and cytoplasmic volume (Mitchison, 2019). We use a bag in a bag model, similar to refs (Deviri and Safran, 2021; Lemière et al., 2021). Following these studies, we assume a spherical cell and nucleus, and that on the obervation time scales both the cell surface and the nuclear surface are at mechanical equilibrium (Salbreux et al., 2007; Venkova et al., 2021). Under these assumptions, a first equation describes the force balance between osmotic pressure and mechanical tension at cell membrane,

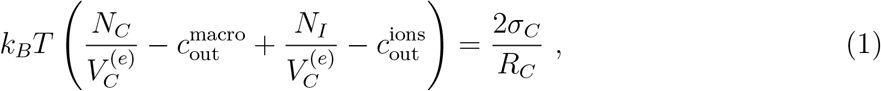

In this equation, the right hand side (ΔΠ) is the osmotic pressure, while the left-hand side (Δ*P*) is the contribution of mechanical forces (in our case, surface tension and the cytoskeleton). In this equation 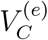 is the accessible cellular volume contributing to osmosis, *R_C_* is the radius of the cell, *N_C_* is the number of osmotically active macromolecules in the cell, while *N_I_* is the number of osmotically active ions in the cell, balancing respectively 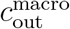 and 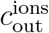 the external concentrations of macromolecules and ions. Finally, *σ_C_* is a mechanical surface tension of the cell. A variant of this equation can consider also the role of osmotically active small osmolytes.

A second equation following the same principles describes mechanical equilibrium between osmotic pressure and mechanical tension at the nuclear envelope. The nucleus has pores, hence we can safely neglect ions and small osmolytes, and assume that only macromolecules are osmotically active (Deviri and Safran, 2021) (see also below)

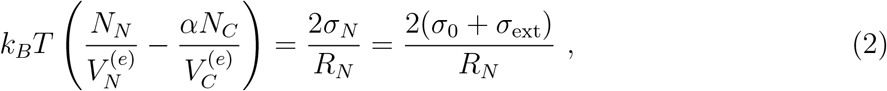

where the notation is similar as Eq. (1), and the suffix *N* stands for nucleus. We note that *N_N_* is the number of osmotically active proteins in the nucleus (roughly, those that have an NLS), equally, we assume that *αN_C_* is the number of osmotically active proteins in the cytoplasm (this is different from *N_C_*, which also includes proteins that can freely shuttle). Finally we have decomposed the tension term into *σ*_0_, a (constitutive) nucleus surface tension and e contribution *σ*_ext_ is a possible from external forces from the cytoskeleton or other active elements, of which we want to test the quantitative effects. Assuming spherical symmetry, *σ*_ext_ is related to external force by the relation *F*_ext_ = 8*πR_N_σ*_ext_ Equation (2) can be seen as a simplified version of the model by Kim *et al*. (Kim et al., 2015), neglecting mechanics of nuclear shape deformations.

A recent study (Andreu et al., 2021) finds that nuclear import increases under force, while export is not biased. Motivated by this, and by our experimental observations, we also propose and examine a model variant incorporating the indirect effects of this force-biased transport on the equilibrium nuclear size. We model this effect in our simple framework as a bias on the nuclear concentration of proteins, based on the external tension, following

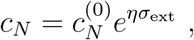

where 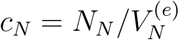 and *η* is a coefficient that quantifies the bias. We estimate the value of *η* directly from experimental data from ref. (Andreu et al., 2021) below.

The main assumption of this model is that cell and nucleus are always in mechanical equilibrium, because the observation time is longer than the relaxation time scales. Additionally, we neglect tension terms for the cell (*σ_C_* ≈ 0), these terms should be small following Deviri and Safran, although they may lead to corrections (Deviri and Safran, 2021). For simplicity, we also always assume spherical symmetry, neglecting any mechanical deformation. These radical assumptions make the model treatable, and should not affect our conclusions, which are based on qualitative behavior or on order-of-magnitude quantitative differences. In standard conditions, external macromolecular osmotic (oncotic) pressure on the cell should be negligible. 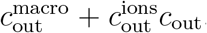, since the hyperosmotic shocks are applied with sucrose, this term becomes relevant in that case (the relevant quantity should be 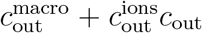. Eq. (2) neglects ions (chromatin counterions) for the nucleus, once again a small contribution according to Deviri and Safran (Deviri and Safran, 2021). Finally, we neglect non-osmotically accessible volume for the nucleus (not measured in our data), 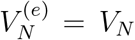 and we assume that for the cell the osmotically accessible volume is the cytoplasmic volume *V_C_*. A variant considering inaccessible volumes as parameters is considered by Deviri and Safran, who show that their predictions are robust. In any case these corrections do not affect qualitative behavior or vary the quantitative behavior by orders of magnitude.

Under these assumptions, the model behavior is simple. External osmotic pressures (and mechano-osmotic coupling from pumps and channels) set a cell volume, and we can envisage a situation where the cell is (essentially) causally uncoupled from the nucleus, and solve Eq. (1) for the cytoplasmic osmotic pressure or equivalently cytoplasmic volume. Then these variables can be used as input variables for the nucleus, solving Eq. (2). While highly simplified - this picture is sufficient to explore different scenarios for our data, as we detail below.

### Estimates of relevant parameters

All the key parameters and/or parameter ranges were fixed from literture values. Osmotic pressures Π_c_ and Π_*n*_. are order 10000 Pa (N/m^2^) (Deviri and Safran, 2021). In order confirm this estimate, we can use the fact that volumes are in the range of 100-1000 *μ*m^3^, while protein concentrations are known to be in the range of 2 – 4 · 10^6^ molecules / *μ*m^3^, which means that in a compartment of ~ 300*μ*m^3^ such as the nucleus there are about 3 · 10^8^ proteins, and in a compartment of ~ 2000*μ*m^3^ such as the cytoplasm there are about 2 · 10^9^ (to which we have to sum the contribution of the small molecules). Below there are some arguments on how many of them should be osmotically active (Milo, 2013). Assuming that a certain number of molecules (probably order 10^8^ – 10^9^ given the above considerations) are osmotically active in a compartment, we can obtain osmotic pressures (Π = *k_B_TN*/*V*). For example for a nucleus of ~ 300*μ*m^3^ with 3 · 10^8^ proteins we get a proessure of 4 kPa. Once again, the estimates are not counting the role of ions and small osmolites in the cytoplasm, but they should be reliable for the nucleus (and a lower bound for the cytoplasm).

Previous studies suggest that about 80% of the proteins are estimated to be localized (either in the nucleus or in the cytoplasm) (Deviri and Safran, 2021). Hence, we estimate that (*αN_C_* + *N_N_*)/(*N_C_* + *N_N_*) = 0.8, which gives *α* ≈ 2/3. Additionally, in yeast, the fraction of *q* of nuclearly-localized to cytoplasmically localized proteins is about 1/2 (Kumar et al., 2002). However, since this factor is a major determinant of the NC volume ratio, and we find ratios that are about 0.2, we conclude that this factor must be smaller in mammalian cells.

The apparent tension of the nucleus, estimating *σ_N_* or *σ*_0_ is around 10^−6^ N/m in nuclear shape fluctuations experiments (Chu et al., 2017; Introini et al., 2021), but this value is an underestimation of the actual tension, since it is affected by excess flickering due to transient deformations of active origin (decreasing the effective tension with respect to the actual mechanical tension). We can assume that this value is a lower bound for the real mechanical tension *σ*_0_.

We assume that cytoskeletal forces are compressive, in line with the recent literature (but we have also considered the opposite case, see below). In order to estimate an upper bound for the external tension generated by these forces, as well as an order of magnitude for cytoskeletal forces, we have considered force traction measurements for cells on substrates of different stiffness. For a substrate stiffness of 5 kPa, we measured a total traction energy of 0.5 fJ from the same cells (Nastaly et al., 2020). With this value, if we assume that all the energy is transmitted on the nucleus (i.e., an upper bound for the tension), that this energy is *σ_N_S*, where *S* is the nuclear surface area. For a nucleus of radius 4 *μm* this gives us *σ_N_* ≃ 10^−5^ N/m. Vianay and coworkers (Vianay et al., 2018) find higher values (about 25 fJ) for RPE1 cells on 40 kPa substrates. This gives *σ_N_* ≃ 5 · 10^−4^ N/m.

We can also consider the total force on a nucleus of radius 4 *μ*m (4 · 10^−6^m), to exert mechanically a pressure of 1000 Pa (this pressure would be able to perturb the osmotic pressure from the nucleus, which as discussed above is of order 4 kPa). Since Δ*P_N_* = 2*σ_N_*/*R_N_* this upper bound would correspond to an (effective) surface tension of about 10^−3^ N/m. This value is considerably higher than the values we estimate from force traction measurements, which can already be considered upper bounds. Since total force is pressure times surface, we would need about 50 nN (50 · 10^−9^N) from the cytoseleton to obtain this force on the whole nuclear surface (still assuming a nucleus of radius 4 *μ*m). This is equivalent to order 50.000 motors exerting pN forces. However, these forces on the nucleus would be observed also by traction force microscopy, by Newton’s third law.

Given these considerations, we have considered a range of 10^−6^ to 10^−3^ N/m for both constitutive and applied mechanical tensions, and we consider that a realistic range for tensions on the nucleus applied by the cytoskeleton could be 10^−5^ – 10^−4^ N/m.

In order to fix the bias parameter in the model variant with force-biased transport, we used the data provided by Andreu and coworkers (Andreu et al., 2021). In this study (Fig 2I of the paper), the authors find a change in the concentration of nuclearly localized proteins *c_N_* by a factor of 3/2 by using gels of stiffness from 1.5 to 30 kPa. Based on the above considerations on force traction measurements, we assume that these conditions could correspond to 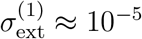 and 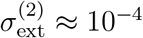 respectively. Assuming as above that

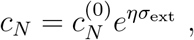

we can now use these two conditions to calibrate this dependency, and find *η*. Since

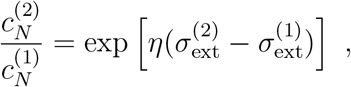

we get *η* ≈ 4 · 10^3^ m/N.

### The bag-in-a-bag model may capture the behavior of hyperosmotic shocks, but is falsified by the experimental observations for cell spreading/detachment

In order to capture the changes of nuclear volume during a hyperosmotic shock, we assume that extracellular ion concentration 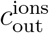 changes, setting a new value for Π_*C*_ = *αN_C_*/(*βV_C_*), with *β* = (*k_B_T*)^−1^. We then solve Eq. (2) for *V_N_* using Π_*C*_ as an input.

**FIG. SM1.**
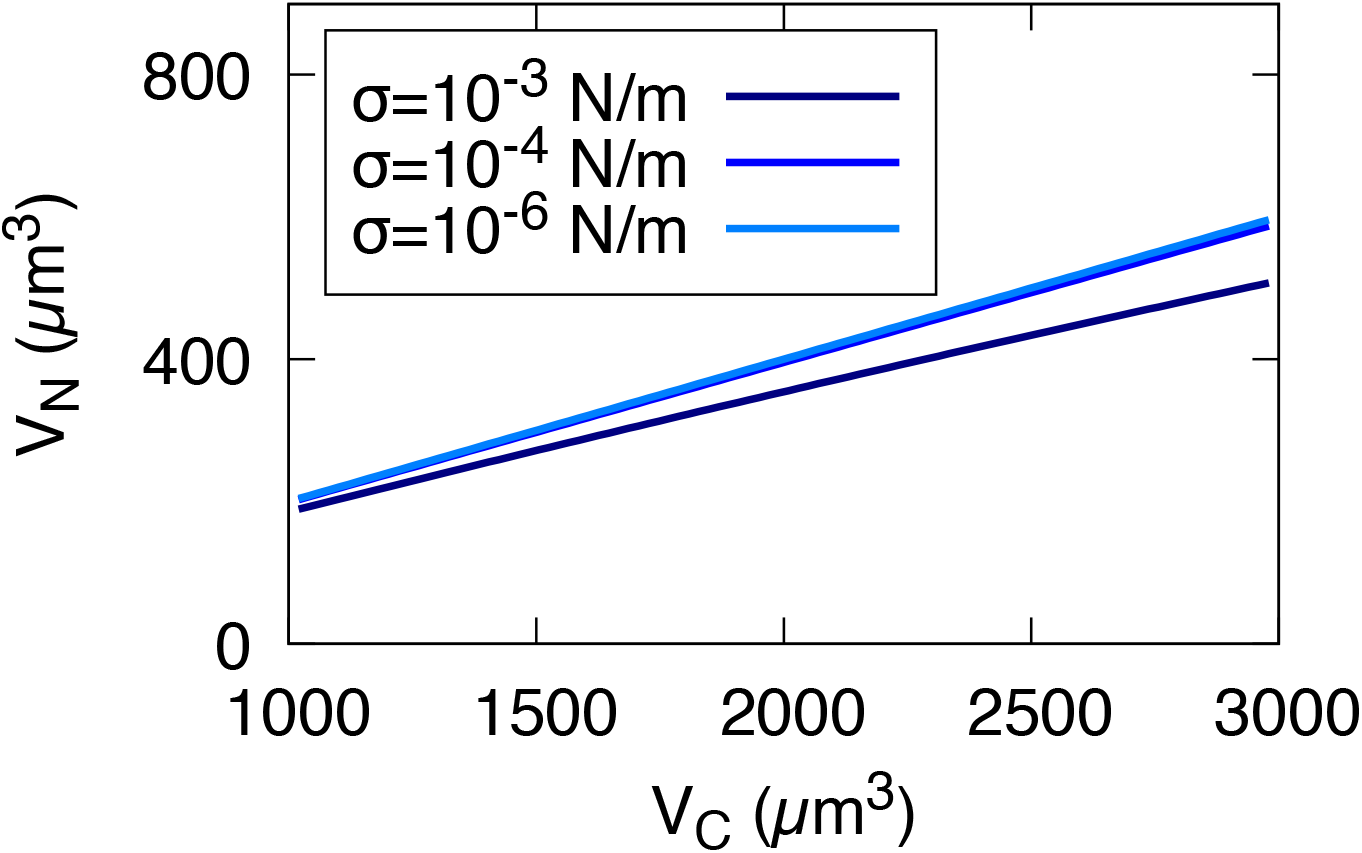
Upon osmotic shocks (on time scales of tens of minutes, before mechano-osmotic regulation takes place (Venkova et al., 2021)), the bag-in-a-bag model predicts nuclear volume to be proportional to cytoplasmic volume, and the proportionality becomes nonlinear only for strong nuclear surface tension. Parameter values: *q* = 0.2 (fraction NLS/NES proteins) *σ_N_* in the range 10^−6^ to 10^−3^ N/m (different solid curves) *N_N_* = 3 · 10^8^ (nuclearly localized macromolecules).

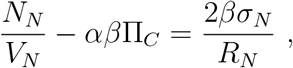

where 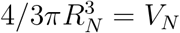.

For negligible *σ_N_*, the solution of this equation is simply

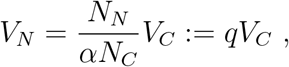

where *q* is the ratio between nuclearly localized (“NLS”) and nuclearly exported (“NES”) proteins, estimated above (from yeast data) to be close to 1/2.

For sufficiently small values of the tension ^1^, we can solve perturbatively the equation and obtain a tension-dependent expression,

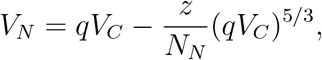

where we have defined *z* = 2*βσ_N_*.

Figure SM1. reports these expressions, for parameter values within the ranges estimated above. The plot shows that (in line with the data) the qualitative model expectation is a proportionality between nuclear volume and cytoplasmic volume, basically set by the ratio of osmotically active particles. Tension, in the range of estimated realistic values, does not affect this trend. It reduces the nuclear volume at fixed cytoplasimc volume, but the changes are only slight. Compressive forces by the cyctoskeleton can contribute to this trend, but again the expected volume reduction is small. We estimate it to be at most less than 10% compared to a tensionless nucleus for the upper bounds of the realistic values for the constitutive tension and the cytoskeletal compressive forces. As a consequence, we find that the nuclear size changes predicted by the bag-in-a-bag model (Deviri and Safran, 2021; Lemière et al., 2021) in presence of changes in external forces are at odds with our experimental observations in cell spreading and detachment experiments. During cell spreading, the cytoskeleton is believed to apply compressive forces to the nucleus, which, would lead to volume-decreasing deformations, in contrast with the observed increase in volume. Viceversa for cell detachment

To sum up, assuming a bag-in-a-bag framework (i) the volume changes observed in osmotic shocks are explained but (ii) the assumption that tension is too small to modify volume in a considerable way is falsified by the data. Crucially, in this prediction the external tension makes a *qualitative* difference. Without an applied external tension, the model is falsified, as it cannot explain why nuclear volume changes during cell spreading, while cytoplasmic volume is roughly constant. However, if the cytoskeleton exterts compressive forces, the model remains at odds with the data, as the nuclear volume increases upon spreading and decreases upon detachment and in suspended cells.

### Force-biased transport can predict the result of spreading/detachment experiments

**FIG. SM2.**
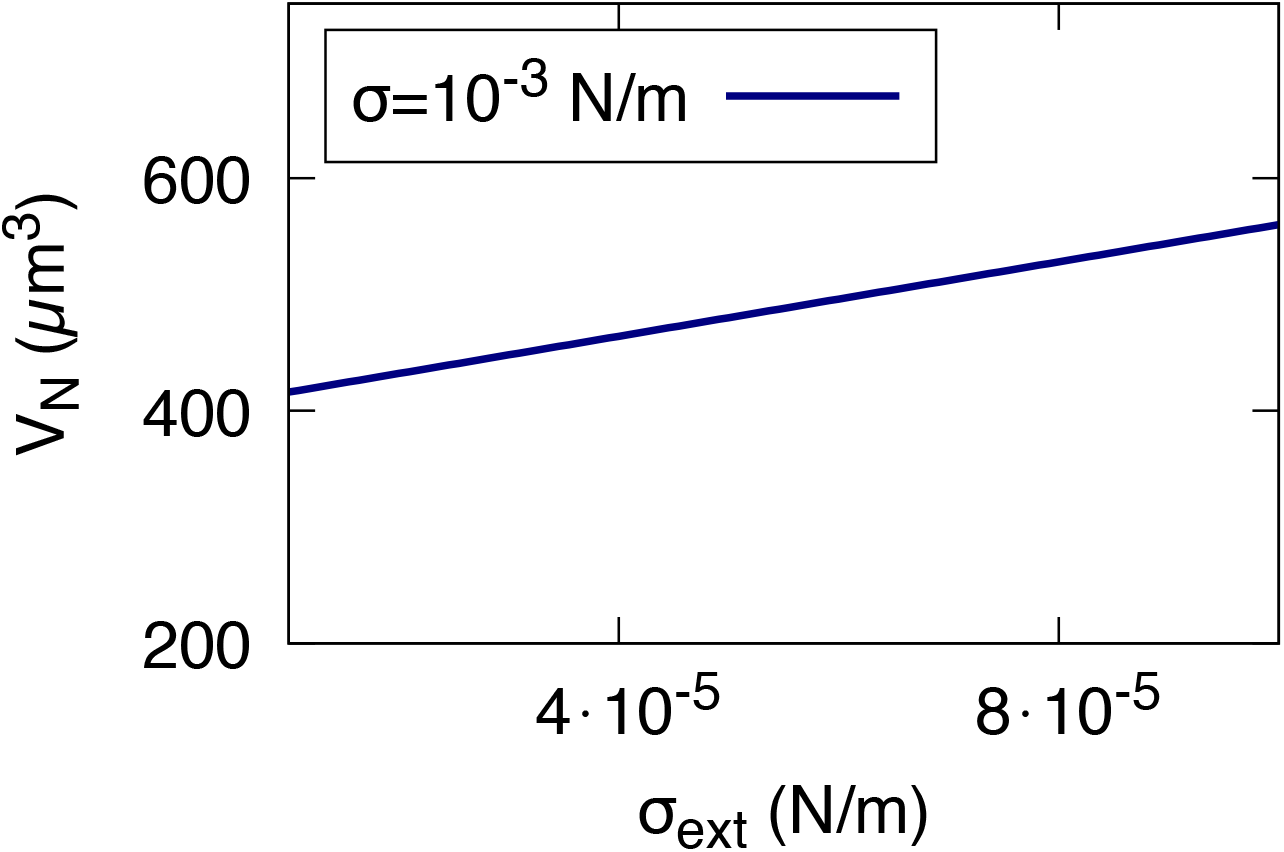
A bias on nuclear import caused by extensile tension can increase nuclear volume more strongly than direct tension, at fixed cytoplasmic osmotic pressure. The plot reports predicted nuclear volume (y axis) for increasing applied extensile tension *σ*_ext_, at fixed basal nuclear surface tension *σ*_0_. Parameter values: *q* = 0.2 (fraction NLS/NES proteins in absence of applied tension) *σ*_0_ = 0 and direct effect of *σ*_ext_ on nuclear size neglected, *η* = 4 · 10^3^ m/N (tension-transport bias) *V_C_* = 2000*μ*m^3^ (cytoplasmic volume).

Conversely, we found that extending the model to the variant including tension-biased transport could realistic values (Fig. 3 in the main text) In the following, we neglect direct effects of surface tension and we assume that only tension-biased transport changes the nuclear volume. This choice is conservative, as well as simplifying the model technically. In this case, nuclear volume is simply computed as

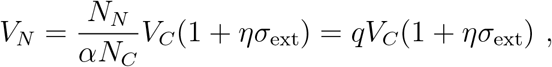

where all the parameters are specified (see above). We can ask how big *η* needs to be, in order to create an increase in nuclear volume of at least 10%. This is given by the condition *ησ*_ext_ ≈ 0.1, i.e. *η* ≈ 1/(10*σ*_ext_).

Our estimate based on data from Andreu and coworkers (Andreu et al., 2021) was that *η* ≈ 4 · 10^3^ m/N. With this value, a tension *σ*_ext_ ≈ 4 · 10^−4^ N/m would be sufficient to achieve visible nuclear deformations. This range of tension is still on the upper end of what we estimate to be achievable from force traction data, but crucially the effect of nuclear deformation on force-biased transport appears to be at least ten times stronger than the direct effect.

We note that we have assumed here that the the impact of the “internal” surface tension on the tension bias is negligible. Technically, this assumption avoids the problem of computing a self-consistent value for the nuclear volume. Biologically, it is possible to speculate that (i) the biased transport saturates at some value of the external forces and/or (ii) tension biased transport may show “adaptation”, in the sense that it is zero in a resting condition (even in presence of some external forces), then responds to external forces perturbing that condition. All these ingredients could be added to the model in a straightforward way. However, given the proof-of-principle aim of our modeling effort, and in absence of precise data to elucidate all these mechanisms, we decided to leave these additional ingredients out of our description. Crucially, the key ingredient of the model is again qualitative: force-biased transport can explain why nuclear volume increases during cell spreading also under the assumption that the cytoskeletal forces felt by the nucleus upon cell spreading are compressive. The model can also explain the qualitative trend of the cytoskeletal perturbations in our experiments, as the release of cytoskeletal compressing forces would remove the force-biased transport, reducing nuclear size.

**FIG. SM3:**
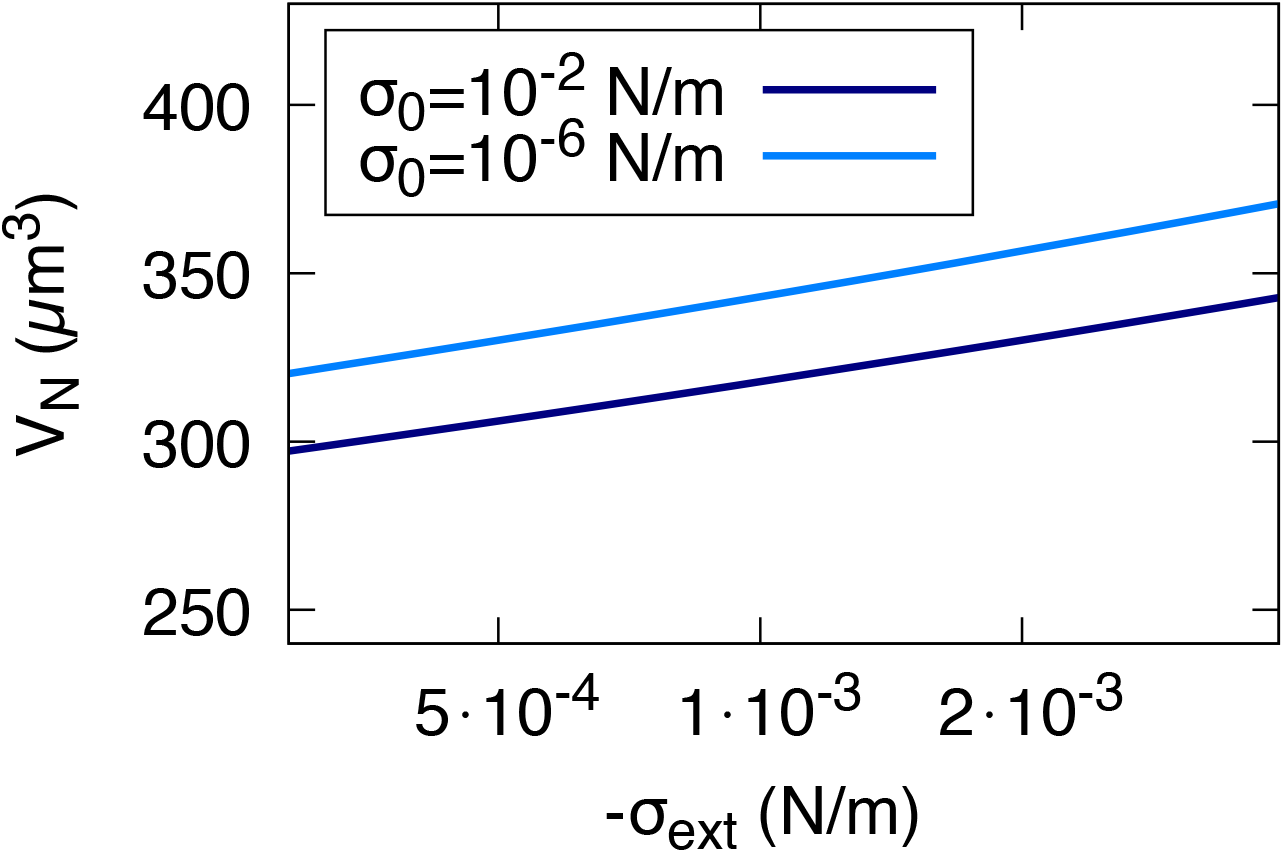
An externally applied extensile tension can mildy increase nuclear volume, at fixed cytoplasmic osmotic pressure. The plot reports predicted nuclear volume (y axis) for increasing applied extensile tension *σ*_ext_, at fixed basal nuclear surface tension *σ*_0_. Parameter values: *α* = 2/3 (fraction of osmotically active proteins that are in the cytoplasm) σ_0_ in the range 10^−6^ to 10^−3^ N/m (different solid curves) Π_*C*_ = 4 kPa (cytoplasmic osmotic pressure).

### The direct effect of extensile forces would also be small

While cytoskeletal forces are generally believed to be compressive, one can also wonder whether cell spreading could associate to extensile forces which directly deform the nucleus, and cell detachment to the release of these forces. To address this question, we also considered the case of *σ*_ext_ < 0, wich in our formalism can describe extensile external forces. In order to explore the effect of an applied (extensile) tension, we solved Eq. (2) taking explicitly into account the external tension *σ*_ext_ and once again for fixed cytoplasmic osmotic pressure Π_*C*_.

In this case it is simple to obtain an implicit expression of *σ*_ext_(*V_N_*) and then get the inverse function by implicit plot,

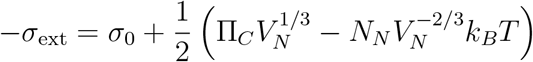

Figure SM3. shows that, according to the bag-in-a-bag model, only (very strong) externally applied forces giving rise to extensile tension of order 10^−3^ – 10^−2^ N/m can noticeably affect nuclear size (in the whole range of possibly relevant constitutive nuclear tensions).

Hence, in spreading experiments (and within the assumptions of our modeling framework), only the presence of an unrealistic increasing extensile force would explain the observed increase of nuclear volume. Hence, even adding a strong extensile tension to the bag-in-a-bag framework leads to the prediction of enormous values of the tension to explain our data.

## Footnotes

1 Specifically, the non-dimensional quantity 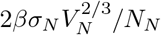 should be small, and in the range of empirically relevant parameter values that we consider this is always the case

